# Sorting endosomes play key roles in presentation of *Mycobacterium tuberculosis*-derived ligands to MAIT cells

**DOI:** 10.64898/2025.12.03.691670

**Authors:** Allison E. Tammen, Jessie C. Peterson, Aneta Worley, David M. Lewinsohn, Elham Karamooz

## Abstract

The immune system has developed specialized mechanisms to recognize intracellular pathogens such as *Mycobacterium tuberculosis* (Mtb). Major Histocompatibility Complex Class I-Related molecule (MR1) is a conserved nonclassical antigen presenting molecule that presents ligands derived from microbial riboflavin synthesis to Mucosal Associated Invariant T (MAIT) cells. While endosomal trafficking facilitates MR1 antigen presentation during Mtb infection, the exact mechanisms by which MR1 loading of Mtb-derived ligands occurs are not known. We found that trafficking through sorting endosomes mediates MR1 antigen presentation during Mtb infection. Sorting endosomes utilize trafficking proteins such as Syntaxin 6, Syntaxin 12, Syntaxin 16 and VAMP4. Prior work demonstrates the importance of VAMP4 for MR1 presentation during Mtb infection; we have found that Stx12 and Stx16 are also important. Interference with Stx12 or Stx16 via siRNA-mediated knockdown reduces MR1 antigen presentation of Mtb. Using RFP-tagged constructs, we found Stx16 co-localized more with MR1 vesicles compared to Stx12 in MR1-GFP expressing airway epithelial cells. Stx12 and Stx16 blockade increase MR1 surface stabilization and total expression, indicating that impaired endosomal trafficking hinders MR1 internalization. Together, these findings support a role for sorting endosomes in the selective sampling of the intracellular environment and MR1-mediated recognition of Mtb-infected cells.

## Introduction

Tuberculosis (TB) caused by the intracellular pathogen *Mycobacterium tuberculosis* (Mtb) is the leading cause of infectious disease morbidity and mortality worldwide^1–4^. Control of Mtb relies on cellular immunity, comprised of both CD4^+^ and CD8^+^ T cells^5–7^. Sampling of the intracellular environment for Mtb is crucial to controlling infection and preventing active tuberculosis^8^. Major Histocompatibility Complex Class I-Related molecule (MR1) is a conserved nonclassical antigen presenting molecule that presents microbial-derived metabolites to Mucosal Associated Invariant T cells (MAITs)^9–12^. MAIT cells, defined by use of the invariant TRAV1-2 T cell receptor, make up approximately 10% of CD8^+^ T cells and 5% of total blood T cells in humans^9,13,14^. They are enriched in mucosal sites like the oral mucosa, digestive tract, skin, airway, and lungs, and have immediate effector function^15–17^. MAIT cells are poised to play a crucial role in controlling Mtb infection, as they are enriched in the lung during Mtb infection, and release important pro-inflammatory cytokines such as IFN-γ in response to Mtb^13,18,19^.

MR1 maintains an intracellular distribution and traffics to the cell surface upon ligand binding^20,21^, suggesting there are tightly controlled mechanisms that regulate the translocation of MR1 to the cell surface. MR1 is consistently found in the endoplasmic reticulum (ER) in the absence of exogenous ligand^21^. After binding of small molecule ligands, such as 5-(2-oxopropylideneamino)-6-d-ribitylaminouracil (5-OP-RU) and 6-formylpterin (6-FP), MR1 associates with Beta-2 microglobulin (β2M), exits the ER, and traffics to the cell surface^21,22^. Additionally, our previous work has shown a proportion of MR1 localizes to endosomal vesicles characterized by expression of Rab7a and Lamp1^20^. To understand mechanisms by which MR1 captures intracellular antigens for presentation to MAIT cells, we screened endosomal trafficking proteins and identified several that play important roles in MR1 presentation during Mtb infection. We demonstrated MR1 requires endosomal trafficking molecules such as Syntaxin 18, Vesicle-Associated Membrane Protein 4 (VAMP4), and Rab6 to activate MAIT cells^20,23^. Importantly, these proteins are not required for HLA-Ia nor HLA-E mediated presentation^20^. We also described different requirements for MR1 presentation of intracellular microbes such as Mtb versus exogenously added antigens^24^.

VAMP4 is a Soluble N-ethylmaleimide sensitive factor (NSF) attachment receptor (SNARE) predominantly localized in the *trans*-Golgi network (TGN) but also present on endosomes. It functions in the endosome-Golgi trafficking pathway to facilitate the sorting and delivery of cargo within the cell^25–27^. Depletion of VAMP4 or its trafficking partners induces Golgi fragmentation, suggesting that VAMP4 containing SNARE complexes are required to maintain the Golgi ribbon structure and maintain normal retrograde trafficking from sorting endosomes to the TGN^25^.

We have demonstrated VAMP4 uniquely contributes to MR1 presentation during Mtb infection, as it is not required for MR1 presentation of 6-FP which is loaded in the ER^20,23^. This would suggest a role for endosome-mediated trafficking of MR1 in presentation of ligands derived from intracellular microbes^20,23^. SNARE proteins such as VAMP4 assemble into complexes to promote membrane fusion events and facilitate trafficking of endosomes to their target locations^25,26^. In eukaryotes, four different SNARE subtypes comprise SNARE complexes^27–29^. VAMP4 (R-SNARE) contains an arginine as the central amino acid residue of the SNARE motif. It partners with Q-SNAREs which contain a glutamine as the central amino acid residue of the SNARE motif. Syntaxins are an important group of Q-SNARE proteins that localize to several intracellular compartments^30–32^. VAMP4 partners with Syntaxin 12 (Stx12) and Syntaxin 16 (Stx16), Qa-SNARES, Vti1a, a Qb-SNARE, and Syntaxin 6 (Stx6), a Qc-SNARE; together, this SNARE complex mediates vesicle docking and membrane fusion events to facilitate sorting endosome to TGN trafficking^27,33,34^.

Localized on sorting endosomes, Stx12 facilitates homotypic fusion to internalize plasma membrane surface proteins^31,35^. Stx12 is also referred to as Syntaxin 13, as they share 98% homology and are orthologs of the same gene^35–38^. Stx16 functions in early endosome to TGN transport and has been shown to be important for retrograde transport of several cargo proteins and cytokinesis of dividing cells^33,34,39–41^. Localized to the Golgi, Stx6 functions in TGN vesicle trafficking and TGN-endosome/endosome-TGN transport^31,42^. The role of sorting endosomes and these VAMP4 trafficking partners they utilize in MR1 presentation during Mtb infection has yet to be fully elucidated. We previously tested the role of endosomal trafficking proteins and found Stx16 knockdown affected MR1 presentation during Mtb infection^24^. While our previous approach established a role for this protein, the goal of this study is to more precisely characterize the role of Stx16 and other VAMP4 trafficking proteins through comprehensive testing across a range of antigen concentrations. Additionally, we sought to characterize the intracellular location of these trafficking proteins relative to MR1 and assess their roles in trafficking of MR1 loaded with exogenous ligand in the ER.

In this study, we evaluate the role of Stx6, Stx12 and Stx16 in MR1-dependent presentation of Mtb in airway epithelial cells. We find Stx12 and Stx16 play key roles in MR1 presentation of Mtb infection, as interference with Stx12 or Stx16 via siRNA-mediated knockdown reduces MR1 antigen presentation of Mtb-derived ligands. We found Stx16 co-localizes with Golgi, and using RFP-tagged constructs, we found Stx16 co-localized more with MR1 vesicles compared to Stx12. Stx12 and Stx16 blockade increase MR1 surface stabilization and total expression, suggesting that impaired endosomal trafficking hinders MR1 internalization. Together, these findings support a unique role for this post-Golgi pathway of MR1 antigen presentation in selective sampling of sorting endosomal compartments for recognition of Mtb-infected cells.

## Results

### Stx12 and Stx16 blockade specifically impair MR1 presentation during Mtb infection

Previously, we conducted a limited analysis of the role of Stx16 in MR1 antigen presentation during Mtb infection^24^. To further characterize the role of Stx16 and establish whether other VAMP4 trafficking partners are also critical for MR1 presentation of Mtb antigens, we performed siRNA inhibition of Stx6, Stx12, and Stx16 in airway epithelial (BEAS-2B) cells. Following infection with Mtb, we tested these cells for their ability to activate a MAIT cell clone. Knockdown of Stx6 reduced *Stx6* transcripts by 80% in 48 hours (Figure 1A). Functionally, compared to a missense control, Stx6 knockdown did not significantly affect MR1 presentation of Mtb antigens to a MAIT cell clone (Figure 1B, p=0.7413). Knockdown of Stx12 reduced *Stx12* transcripts by 80% in 48 hours (Figure 1C). Compared to a missense control, Stx12 knockdown significantly reduced MR1 presentation of Mtb antigens to a MAIT cell clone, decreasing the max response from an average of 664±56 to 455±115 IFN-γ Spot Forming Units (SFUs), a 31% decrease (Figure 1D, p=0.0333). To determine the specificity of Stx12 knockdown on MR1 antigen presentation, we also tested the effect of Stx12 knockdown on HLA-B45 antigen presentation using an HLA-B45-restricted T cell clone specific for an epitope derived from Mtb CFP10. We found that Stx12 knockdown had no significant effect on HLA-B45 presentation of Mtb to an HLA-B45-restricted T cell clone (Figure 1E, p=0.3375), showing that Stx12 knockdown does not alter antigen uptake, intracellular infectivity, or cause cellular toxicity. Knockdown of Stx16 reduced *Stx16* transcripts by 74% in 48 hours (Figure 1F). Similarly to Stx12 knockdown, Stx16 knockdown significantly reduced MR1 presentation of Mtb antigens to a MAIT cell clone, decreasing the max response from an average of 716±37 to 393±58 SFU, a 45% decrease (Figure 1G, p<0.0001), but had no significant effect on HLA-B45 presentation of Mtb to an HLA-B45-restricted T cell clone (Figure 1H, p=0.2345). Together, these data indicate that Stx12 and Stx16, but not Stx6, play a role in presentation of Mtb to MAIT cells by MR1.

**Figure 1:**
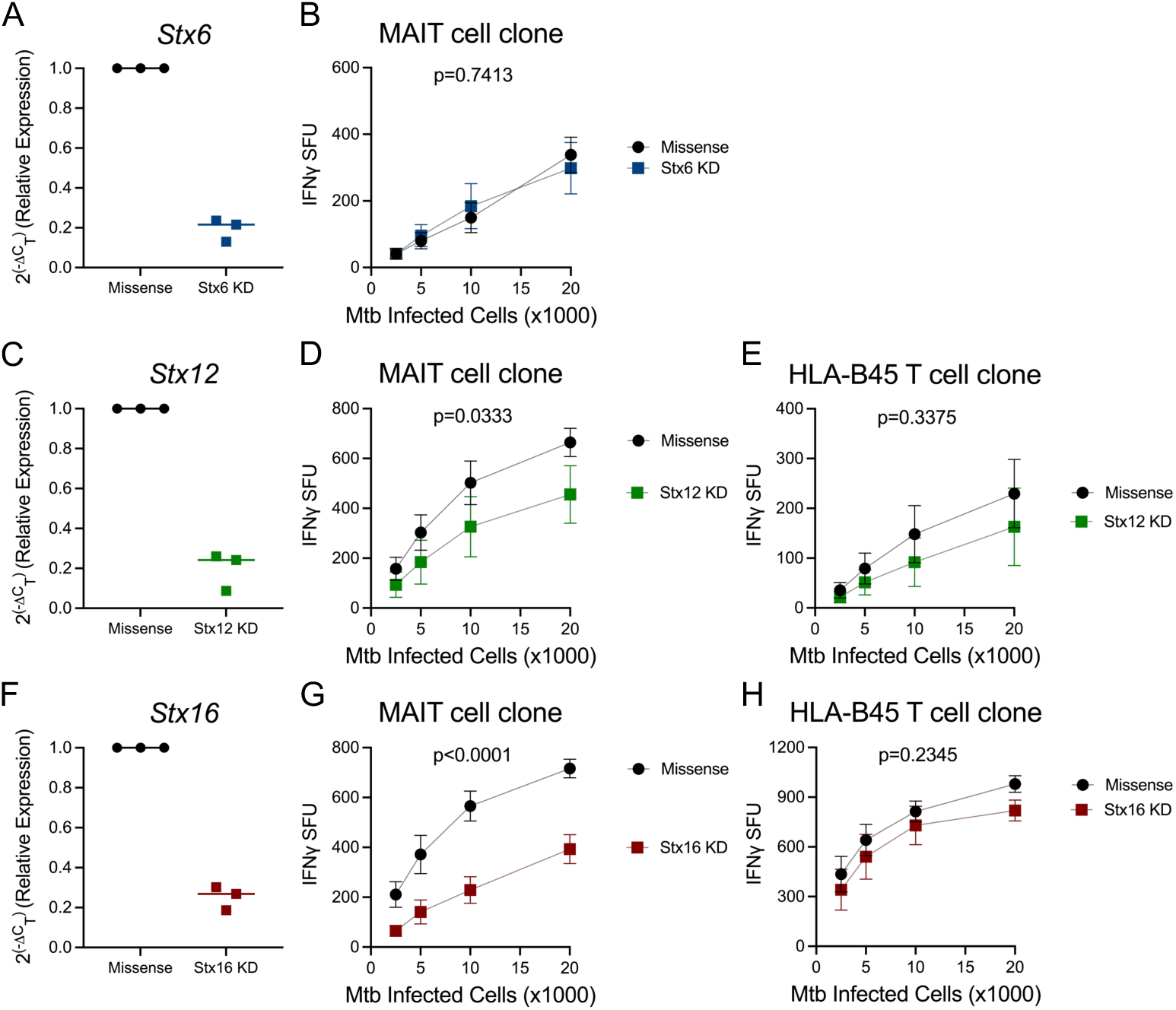
Stx12 and Stx16 blockade specifically impair MR1 presentation during Mtb infection. A) RT-qPCR of RNA isolated from Missense control and Stx6 knockdown (KD) BEAS-2Bs. *Stx6* expression was calculated relative to *Gapdh.* B) IFN-γ ELISpot assay with Mtb infected Missense control and Stx6 knockdown BEAS-2Bs using MR1-restricted MAIT cell clone, p=0.7413. MAIT cell responses measured by mean IFN-γ spot forming units (SFU). C) RT-qPCR of RNA isolated from Missense control and Stx12 knockdown BEAS-2Bs. *Stx12* expression was calculated relative to *Gapdh*. D) IFN-γ ELISpot mean SFU from Mtb infected Missense control and Stx12 knockdown BEAS-2Bs using MR1-restricted MAIT cell clone, p=0.0333. E) IFN-γ ELISpot mean SFU from Mtb infected Missense control and Stx12 knockdown BEAS-2Bs using HLA-B45-restricted T cell clone D466-D6, p=0.3375. F) RT-qPCR of RNA isolated from Missense control and Stx16 knockdown BEAS-2Bs. *Stx16* expression was calculated relative to *Gapdh*. G) IFN-γ ELISpot mean SFU from Mtb infected Missense control and Stx16 knockdown BEAS-2Bs using MR1-restricted MAIT cell clone, p<0.0001. H) IFN-γ ELISpot mean SFU from Mtb infected Missense control and Stx16 knockdown BEAS-2Bs using HLA-B45-restricted T cell clone D466-A10, p=0.2345. ELISpot assay data are pooled from n=3 independent experiments. Error bars represent standard error of the mean (SEM). Top and EC_50_ SFU values were determined by least-squares regression and used to calculate p-values by extra sum-of-squares F test.

### Stx16 localizes to the TGN

Stx16 is a tail-anchored membrane protein localized mainly at the TGN^33,43^. To confirm the location of Stx16 in airway epithelial cells, we performed immunofluorescence staining in BEAS-2Bs. We also stained for the TGN membrane using an antibody against golgin-97. We found that Stx16 staining overlaps with golgin-97 (Figure 2A). We next wanted to observe how Stx16 knockdown affects the distribution of the TGN and performed Stx16 and golgin-97 immunofluorescence staining in BEAS-2Bs 48 hours after siRNA inhibition of Stx16. Stx16 knockdown resulted in a substantial decrease in Stx16 staining (Figures 2B, 2C). Additionally, Stx16 knockdown resulted in dispersion of the Golgi, but the overall intensity of golgin-97 was unchanged (Figure 2B, 2D). This dispersion of the Golgi is consistent with previous work showing that Stx16 and other TGN SNAREs like VAMP4 are required to maintain the ribbon structure of the TGN^25^. While it is known Stx12 localizes to sorting endosomes and the tubular vesicle network associated with sorting endosomes^27,32^, we were unable to find a suitable Stx12 antibody to confirm this localization pattern in BEAS-2Bs.

**Figure 2:**
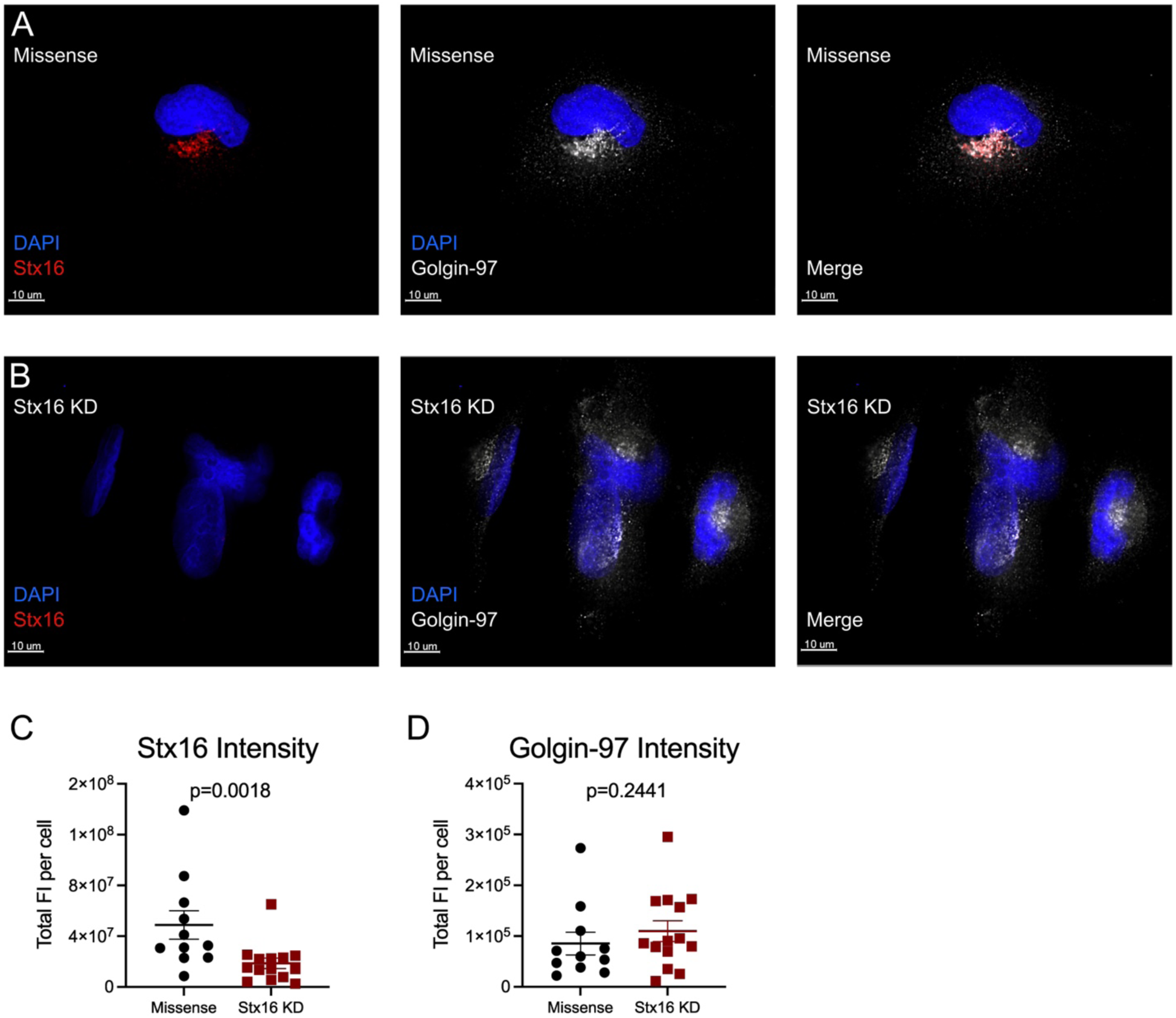
Stx16 localizes with the TGN. A) Missense control and B) Stx16 knockdown (KD) BEAS-2B:TET-MR1GFP LV cells stained with antibodies against Stx16 (left) and golgin-97 (center); merge of Stx16 and golgin-97 (right). Representative images from n=2 independent experiments. A single Z-stack is shown. Scale bars represent 10µm. C-D) Sum fluorescence intensity (FI) per cell values from n=2 independent experiments. C) Total FI per cell values of Stx16 staining, p=0.0018. D) Total FI per cell values of golgin-97 staining, p=0.2441. Statistical significance between Missense control and Stx16 knockdown determined using Mann-Whitney test for nonparametric data.

### Stx12 and Stx16 co-localize with MR1 vesicles

Next, we sought to characterize the interaction between MR1 and Stx12 and Stx16. To do this, we created Stx12 and Stx16 constructs tagged with red fluorescent protein (Stx12-RFP and Stx16-RFP). We first transiently transfected Stx16-RFP into BEAS-2Bs stably transduced with a doxycycline inducible MR1-GFP construct (BEAS-2B:TET-MR1GFP LV)^44,23^ and Stx16 protein expression was evaluated by western blot using a Stx16 antibody. Western blot analysis showed staining for both native Stx16 and Stx16-RFP compared to non-target control transfected cells (Figure 3A). To measure the association between MR1 and Stx12 and Stx16, BEAS-2B:TET-MR1GFP LV cells were transiently transfected with the RFP-tagged constructs, treated with doxycycline to induce MR1-GFP synthesis, and imaged via live-cell fluorescence microscopy the following day. We found 51% (min 20%, max 74%) of MR1 vesicles co-localize with Stx16 while 31% (min 10%, max 53%) co-localize with Stx12 (Figure 3B). The degree of co-localization between Stx16 and MR1 compared to Stx12 and MR1 is significantly higher (Figure 3B, p<0.0001). These findings indicate that while both Stx12 and Stx16 are found in close proximity to MR1 vesicles, suggesting roles during MR1 trafficking, Stx16 exhibits a higher degree of co-localization with MR1.

**Figure 3:**
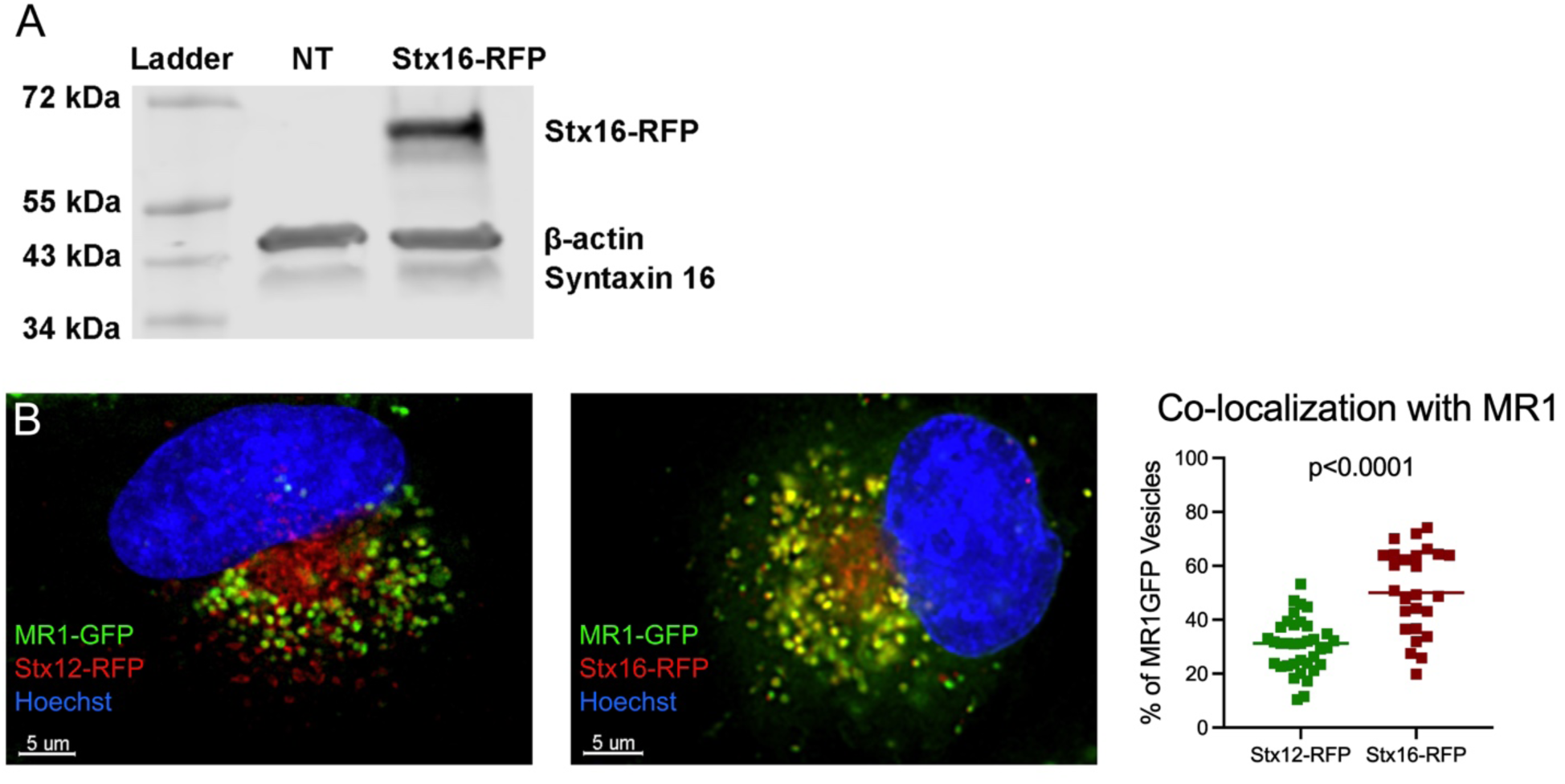
Stx12 and Stx16 co-localize with MR1 vesicles. A) Western blot of Stx16, beta actin, and Stx16-RFP in non-transfected control (NT) and Stx16-RFP transfected BEAS-2B:TET-MR1GFP LV cells. Representative of n=2 independent experiments. B) Stx12-RFP (left) and Stx16-RFP (center) transfected BEAS-2B:TET-MR1GFP LV incubated overnight with doxycycline. Merge of Hoechst, MR1-GFP, and Stx12-RFP or 16-RFP. Representative images from at least n=2 independent experiments. A single Z-stack is shown. Scale bars represent 5µm. Percentage of MR1-GFP Vesicles that co-localize with Stx12-RFP or Stx16-RFP (right). Statistical significance between Stx12-MR1 co-localization and Stx16-MR1 co-localization determined using Mann-Whitney test for nonparametric data, p<0.0001.

### Stx12 and Stx16 blockade increase MR1 surface stabilization and total expression

To further determine how Stx12 and Stx16 affect MR1 antigen presentation, we performed flow cytometry to characterize the effects of Stx12 and Stx16 knockdown in BEAS-2Bs constitutively over-expressing MR1-GFP under a minimal CMV promoter^20^ (MR1-GFP BEAS-2Bs) on MR1 or HLA-A, B, and C cell surface translocation. The small molecule 6-FP is an MR1 antagonist derived from the photodegradation of folic acid^12^. Although unable to activate MAITs, 6-FP is a potent MR1 ligand that is loaded in the ER and induces MR1 translocation to the cell surface^10,21,45^. We found Stx12 knockdown significantly increased MR1 surface stabilization compared to Missense control with NaOH solvent control treatment (Figure 4A, p=0.0020) and with 6-FP treatment (Figure 4A, p=0.0008). Additionally, we found Stx12 knockdown significantly decreased HLA-A, B, and C cell surface stabilization compared to Missense control with NaOH solvent control treatment (Figure 4B, p=0.0273) but not with 6-FP treatment (Figure 4B, p=0.5967). The effects on HLA-A, B, and C cell surface stabilization with NaOH solvent control treatment were statistically significant but appear more modest; these data and our observation that Stx12 does not significantly affect HLA-B45 presentation of intracellular Mtb (Figure 1E) support the notion that Stx12 knockdown specifically affects MR1 trafficking and presentation. We also found total MR1 expression, as measured by GFP gMFI, to be significantly increased in Stx12 knockdown MR1-GFP BEAS-2Bs compared to Missense control with NaOH solvent control treatment (Figure 4C, p=0.0068) and with 6-FP treatment (Figure 4C, p=0.0014). This would suggest impaired internalization of either endocytosed MR1 or in endocytosis of MR1 following surface translocation with Stx12 knockdown.

**Figure 4:**
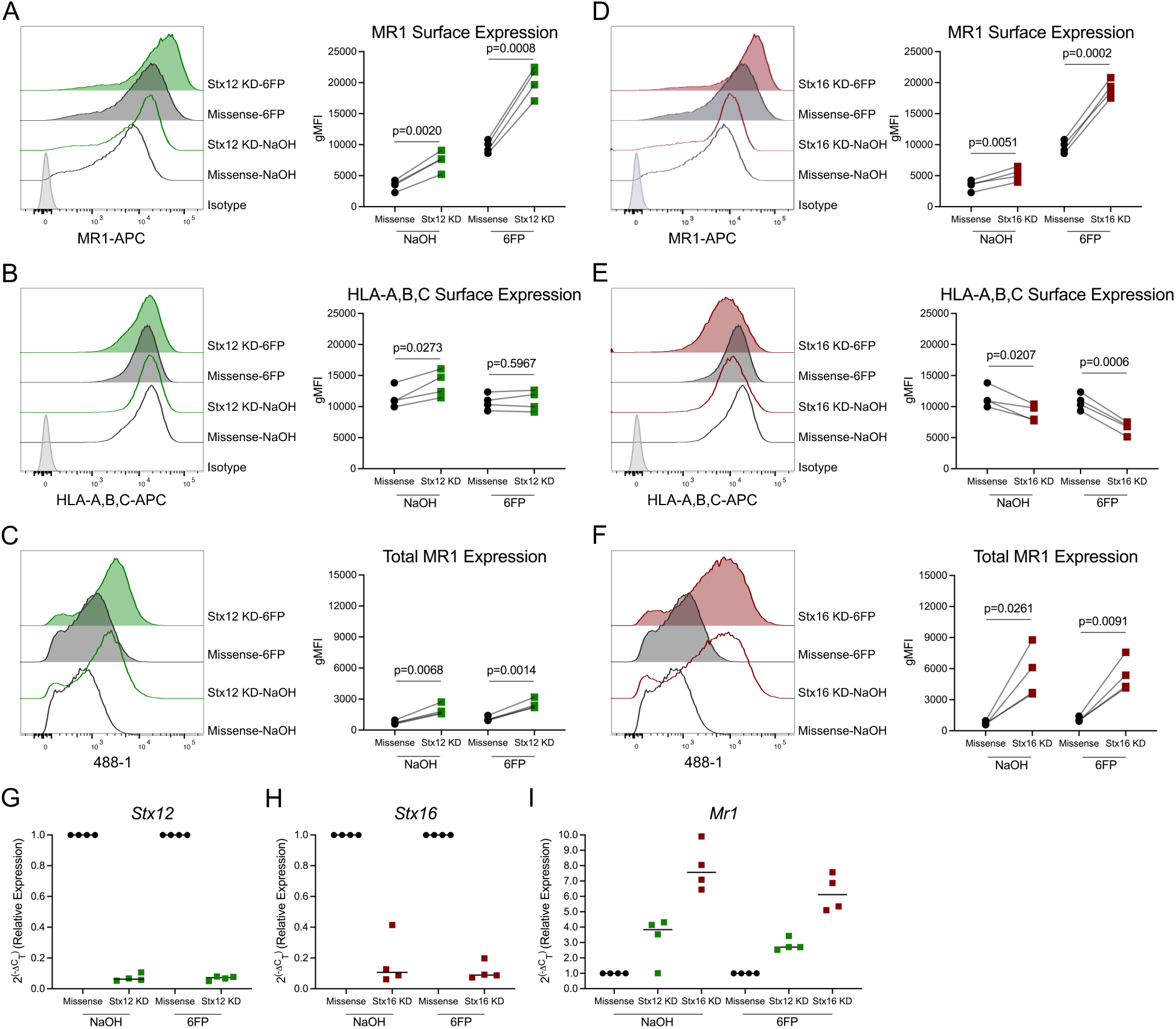
Stx12 and Stx16 blockade increase MR1 surface stabilization and total expression. A-F) Flow cytometry analyses of 100µM 6-FP or 0.01M NaOH solvent control treated Missense control, Stx12, or Stx16 knockdown (KD) MR1-GFP BEAS-2Bs. Representative histograms (left) and gMFI values (right). A) MR1 surface expression in Missense control and Stx12 knockdown MR1-GFP BEAS-2Bs treated with NaOH solvent control, p=0.0051, or treated with 6-FP, p=0.0002. B) HLA-A,B,C surface expression in Missense control and Stx12 knockdown MR1-GFP BEAS-2Bs treated with NaOH solvent control, p=0.0207, or treated with 6-FP, p=0.0006. C) Total MR1 expression (GFP gMFI) in Missense control and Stx12 knockdown MR1-GFP BEAS-2Bs treated with NaOH solvent control, p=0.0261, or treated with 6-FP, p=0.0091. D) MR1 surface expression in Missense control and Stx16 knockdown MR1-GFP BEAS-2Bs treated with NaOH solvent control, p=0.0020, or treated with 6-FP, p=0.0008. E) HLA-A,B,C surface expression in Missense control and Stx16 knockdown MR1-GFP BEAS-2Bs treated with NaOH solvent control, p=0.0273, or treated with 6-FP, p=0.5967. F) Total MR1 expression (GFP gMFI) in Missense control and Stx16 knockdown MR1-GFP BEAS-2Bs treated with NaOH solvent control, p=0.0068, or treated with 6-FP, p=0.0014. G-I) RT-qPCR of RNA isolated from Missense control, Stx12, and Stx16 knockdown MR1-GFP BEAS-2Bs. *Stx12*, *Stx16,* and *MR1* gene expression were calculated relative to *Gapdh*. Flow cytometry data are pooled from n=4 independent experiments. gMFI values were used to calculate p-values by two-tailed paired t-test.

We found Stx16 knockdown significantly increased MR1 surface stabilization compared to Missense control with NaOH solvent control treatment (Figure 4D, p=0.0051) and with 6-FP treatment (Figure 4D, p=0.0002). Additionally, we found Stx16 knockdown significantly decreased HLA-A, B, and C cell surface stabilization compared to Missense control with NaOH solvent control treatment (Figure 4E, p=0.0207) and with 6-FP treatment (Figure 4E, p=0.0006). These results would suggest that Stx16 knockdown has a broad spectrum of activity; however we did not observe Stx12 knockdown significantly affecting HLA-B45 presentation of intracellular Mtb (Figure 1H). We also found total MR1 expression, as measured by GFP gMFI, to be significantly increased in Stx16 knockdown MR1-GFP BEAS-2Bs compared to Missense control with NaOH solvent control treatment (Figure 4F, p=0.0261) and with 6-FP treatment (Figure 4F, p=0.0091).

We confirmed percentage knockdowns of Stx12 and Stx16 by RT-qPCR; knockdown of Stx12 reduced *Stx12* transcripts by 93% in NaOH treated cells and by 93% in 6-FP treated cells (Figure 4G). Knockdown of Stx16 reduced *Stx16* transcripts by 83% in NaOH treated cells and by 89% in 6-FP treated cells (Figure 4H). We also compared MR1 expression in Stx12 and Stx16 knockdown BEAS-2Bs compared to Missense control. Knockdown of Stx12 increased *Mr1* transcripts by 3.3-fold in NaOH treated cells and by 2.8-fold in 6-FP treated cells (Figure 4I). Knockdown of Stx16 increased *MR1* transcripts to a greater extent than Stx12 knockdown, raising *Mr1* transcripts by 7.9-fold in NaOH treated cells and by 6.2-fold in 6-FP treated cells (Figure 4I). This would suggest that Stx12 and Stx16 may play roles in regulating vesicular trafficking events that indirectly affect MR1 transcriptional responses. Together, these data indicate that Stx12 and Stx16 inhibition increase MR1 surface expression with or without 6-FP treatment, increase total MR1 expression, and increase *Mr1* mRNA transcripts.

## Discussion

MR1 is a fundamentally novel antigen presenting molecule in that it presents microbial metabolites. Given that very little MR1 is present on the cell surface under basal conditions, and that MR1 only traffics to the cell surface after it has captured an antigen, trafficking of MR1 is likely a tightly regulated process. Elucidating these mechanisms is important for building a more complete understanding of how MR1 trafficking and antigen presentation are regulated, but also how the immune system samples the intracellular environment for ligands that are in minute quantities. It has been well characterized that soluble, exogenous sources of MR1 ligands like 6-FP are loaded on ER-resident MR1, thus inducing MR1 cell surface translocation^12,21,45^. However, there are also distinct pathways that govern MR1 presentation of intracellular pathogens such as Mtb.

We have previously demonstrated that Mtb-infected epithelial cells are capable of stimulating IFN-γ production by MAIT cells^18,46^. Utilizing Mtb deletion mutants of riboflavin metabolism genes, we have shown that deletion of early pathway *Rib* genes that lead to the production of 5-amino-6-D-ribitylaminouracil (5-A-RU) profoundly reduced MAIT cell activation by Mtb^47^. These data support a central role for the riboflavin pathway, specifically 5-A-RU, in the production of Mtb-derived MR1 ligands that can be recognized by MAIT cells. While these Mtb-derived ligands could be loaded on ER-resident MR1, our work supports a unique role for distinct post-Golgi pathways involving endosomal trafficking^20,23^. Our previous work also demonstrated that following cell surface translocation, 6-FP-loaded MR1 is recycled to a compartment where it is optimally available for loading of exogenously added ligands^24^. It has been suggested that MR1 recycling is not required for exchange of exogenously added ligands, as cells deficient in AP2 complex subunit α1 (AP2A1), required for MR1 internalization from the plasma membrane, were able to exchange 6-FP for MAgA-TAMRA, an epitope-tagged derivative MR1 antigen analog tethered to tetramethylrhodamine fluorophore^22,48^. However, work from our laboratory supports that exchange of exogenously added ligands does require internalization, and ligand transfer occurs in post-ER compartments^49^. We have shown that intracellular exchange is required for the presentation of MR1 monomer-delivered 5-OP-RU, requiring the transfer of ligand from the soluble molecule onto endogenous host cell MR1 in Rab5 or LAMP1 labeled compartments^49^. Together, these data support the concept that there are multiple non-redundant pathways by which MR1 samples ligands derived from both extracellular and intracellular environments.

In this study, we expand upon our previous work identifying different endosomal trafficking proteins involved in MR1 presentation specifically during Mtb infection. Although the ER has been identified as the canonical loading pathway for exogenously delivered antigens, our work in airway epithelial cells and Mtb has identified the endosome-Golgi pathway as a key pathway, demonstrated by the importance of VAMP4 and Rab6 in MR1-dependent presentation of Mtb antigens^20,23^. This contrasts other previous work that identified Syntaxin 4 to play a role in MR1 presentation of exogenous antigens only, with no role in MR1 presentation during Mtb infection^24^. We also previously established an effect of Stx16 knockdown on MR1 presentation during Mtb infection^24^. Here, we more quantitatively expand upon those data and characterize other VAMP4 trafficking partners. We find Stx6 knockdown had no role in MR1 antigen presentation during Mtb infection, whereas Stx12 and Stx16 knockdowns had significant effects on MR1 presentation.

Stx12 and Stx16 are Qa-SNAREs, which possess N-terminal antiparallel three-helix bundles compared to Qb- and Qc-SNAREs.^29^ The fact that various Qb- or Qc-SNAREs can comprise the t-SNARE assembly would indicate that Qc-SNARE Stx6 may be dispensable for t-SNARE assembly and association with v-SNARE VAMP4 in this context^50^. This would support our data that Stx6 knockdown does not affect MR1 presentation during Mtb infection. While Vti1A is another VAMP4 trafficking partner, it is a v-SNARE on vesicular membranes that largely functions in intra-Golgi traffic in mammalian cells^38,51^. Thus, we focused our efforts on Stx6, Stx12, and Stx16, as these are endosomal membrane t-SNAREs that interact with VAMP4.

We evaluated the intracellular expression of Stx16 in BEAS-2B airway epithelial cells and found Stx16 co-localized with golgin-97, a marker of the TGN membrane. Stx16 knockdown did not affect the intensity of golgin-97 by microscopy but resulted in dispersion of the Golgi as visually apparent in Figure 2. As TGN SNARES like VAMP4 and Stx16 are required to maintain the ribbon structure of the TGN^25^, impaired VAMP4-SNARE complex formation likely causes impaired endosome to TGN trafficking, thus destabilizing the TGN organization and leading to dispersion. Although we were unable to find a suitable Stx12 antibody, previous work has demonstrated Stx12 co-labels with Rab5 and localizes within the tubular structures of sorting endosomes^27,32^. Despite not localizing to the TGN like its SNARE trafficking partners VAMP4 and Stx16, Stx12 participates in mediating the homotypic fusion of early endosomes.

Via live-cell imaging, we observed MR1 vesicles co-localize with both Stx12 and Stx16 RFP-tagged constructs. This would support the notion that MR1 resides in non-ER, endosomal compartments as demonstrated in our previous work^20,23^, and that VAMP4 trafficking partners are also in very close proximity to or in the same endosomal compartments. Additionally, we quantified the percentage of co-localization between Stx12 or Stx16 and MR1 and found a statistically significantly higher degree of co-localization between Stx16 and MR1 compared to Stx12 and MR1. While MR1 exhibits less co-localization with Stx12 compared to Stx16, we still demonstrate a role for Stx12 in MR1 presentation during Mtb infection. A definitive explanation of this difference between Stx12 and Stx16 co-localization with MR1 and how it may affect MR1 trafficking has yet to be elucidated.

This work expands upon our identification of endosomal trafficking proteins that play a role in MR1 presentation during Mtb infection, as the knockdowns of Rab6, VAMP4, Stx12, Stx16, and Stx18 significantly impair MR1-dependent MAIT cell activation (Table 1)^20,23,24^. To better understand whether the effects of Stx12 and Stx16 knockdowns we observe on MR1 antigen presentation are due to impaired MR1 trafficking or differing quantities of MR1, we measured cell surface translocation and total expression of 6-FP-loaded MR1. Interestingly, the knockdowns of these proteins have variable effects on cell surface translocation of 6-FP-loaded MR1 (Table 1). Unlike Rab6 and VAMP4, where knockdown has no effect on MR1 surface translocation, nor Stx18, where knockdown decreased MR1 surface translocation, we find that Stx12 and Stx16 knockdown significantly increase both surface and total MR1 expression. There are multiple mechanisms that may explain this observation of increased abundance of MR1 alongside reduced antigen presentation function. Firstly, Stx12 and Stx16 may play a role in MR1 internalization through the endocytic pathway. This aligns with previous results that Rab6 inhibition also increased total MR1 expression^23^, suggesting that knockdown of Stx12 or Stx16 impairs MR1 internalization. Alternatively, Stx12 and Stx16 may regulate MR1 trafficking in the context of surface translocation, suggesting Stx12 or Stx16 knockdown reduces VAMP4-SNARE complex formations and affects the ability of MR1 to traffic through endosomal compartments.

**Table 1:** Summary of endosomal trafficking proteins and their roles in MR1 presentation during Mtb infection and 6-FP dependent surface translocation of MR1.

|  | Role in endosomal trafficking: | Effect of knockdown on MR1 presentation during Mtb infection? | Effect of knockdown on cell surface translocation of 6-FP-loaded MR1? |
| --- | --- | --- | --- |
| Rab6 | GTPase | ↓ | ↔ |
| VAMP4 | R-SNARE | ↓ | ↔ |
| Syntaxin 12 | Qa-SNARE | ↓ | ↑ |
| Syntaxin 16 | Qa-SNARE | ↓ | ↑ |
| Syntaxin 18 | Qa-SNARE | ↓ | ↓ |

We also find that mRNA transcripts of MR1 are substantially increased with Stx12 or Stx16 knockdown, indicating that there is a transcriptional effect on MR1. There are multiple hypotheses that could support this finding. Stx12 and Stx16 may facilitate the trafficking of a transcription factor that initiates more *Mr1* transcription, or stabilize *Mr1* transcripts. There is an incomplete understanding of MR1 transcriptional regulation. While much is unknown, it was recently demonstrated IFN-γ stimulates Interferon regulatory factor 1 (IRF1) expression in airway epithelial cells, leading to IRF1 binding to the MR1 promoter and induction of *Mr1* transcription^52^. Additionally, dysregulation of the Ras/ERK pathway has been found to upregulate *Mr1* transcription. Inhibition of ERK1/2 kinases has been shown to increase *Mr1* transcription and protein expression, and these increases were mediated by the transcription factor E74-like ETS transcription factor 1 (ELF1)^53^. In this study, our data could support a model where Stx12 and/or Stx16 inhibition and impaired endosomal trafficking induces IRF1, ELF1, or another unknown transcription factor to upregulate *Mr1* transcription. Alternatively, Stx12 or Stx16 inhibition could stabilize *Mr1* transcripts, or elevated *Mr1* transcripts could be an artifact of constitutive over-expression of MR1 in the MR1-GFP BEAS-2Bs. Given that endogenous MR1 is weakly expressed on the cell surface in WT BEAS-2Bs, we utilized MR1 over-expressing cells to evaluate effects on MR1 cell surface stabilization; this could possibly account for the observed increases in *Mr1* transcripts. These proposed mechanisms require further investigation.

In conclusion, we have identified several endosomal trafficking proteins required for MR1 antigen presentation during intracellular Mtb infection. Importantly, knockdown of these proteins does not affect HLA-IA presentation, indicating this pathway is specific to MR1. Our data is consistent with a model where MR1 resides in non-ER vesicular compartments that utilize Stx12 and Stx16 for trafficking and endocytosis. During intracellular Mtb infection, this permits MR1 to access ligands derived from Mtb in the endosomal trafficking pathway for subsequent loading, trafficking to the plasma membrane, and MAIT cell activation.

## Materials and Methods

### Human Subjects

This study was conducted according to the principles expressed in the Declaration of Helsinki. Study participants, protocols, and consent forms were approved by the Institutional Review Board at Oregon Health & Science University (OHSU) (IRB00000186). All ethical regulations relevant to human research participants were followed. Written and informed consent was obtained from all donors. Peripheral blood mononuclear cells (PBMC) and human serum from human subjects were used to expand T cell clones and in ELISpot medium as described in the methods section.

### Bacteria and cells

*Mycobacterium tuberculosis* (Mtb) H37Rv (ATCC) was grown in Middlebrook 7H9 broth supplemented with Middlebrook ADC, 0.05% Tween-80, and 0.5% glycerol. The bacteria were passaged 10–20 times through a 27-gauge tuberculin syringe before infection. Multiplicity of infection of 8 was used for all Mtb experiments. All experiments with Mtb were done in a Biosafety Level 3 laboratory. Waste was decontaminated in 3% Wescodyne and autoclaved. All other experiments were performed in a Biosafety Level 2 laboratory.

BEAS-2B (ATCC), BEAS-2B constitutively over-expressing MR1-GFP under a minimal CMV promoter^20^, and BEAS-2B transduced with a lentiviral construct encoding tetracycline-inducible, GFP-tagged MR1A (TET-MR1GFP LV)^44,23^ were cultured in DMEM (Gibco) supplemented with 2% L-glutamine (Gibco) and 10% heat inactivated fetal bovine serum (FBS, Gemini).

The following human T cell clones were used: TRAV1–2+ MR1-restricted (D426-G11)^14^, HLA-B45-restricted (D466-A10, minimal epitope CFP10_2–9_)^54^, and HLA-B45-restricted (D466-D6, minimal epitope CFP10_2-12_)^54^.

### Reagents and antibodies

Phytohemagglutinin (PHA, Sigma-Aldrich) was resuspended to 10mg/mL in 10% human serum in RPMI (Gibco) supplemented with L-glutamine and gentamicin. Paraformaldehyde (Electron Microscopy Sciences) was obtained as a 16% stock and diluted to a 4% working stock using phosphate buffered saline (Corning). FBS (GeminiBio) was heat inactivated at 56°C for 45 minutes. Human serum was obtained from donors and heat inactivated at 56°C for 45 minutes. Golgin-97 mouse monoclonal antibody (CDF4, ThermoFisher) was resuspended at a concentration of 1.0 ug/µL and used at 1:100. Syntaxin 16 rabbit monoclonal antibody (ERP9156, Abcam) was used at 1:100. Silencer Select siRNAs (ThermoFisher) were resuspended at concentrations of 50 µM: Silencer Select Negative Control #1 (catalog #4390843), Syntaxin 6 (S19958), Syntaxin 12 (S24307), and Syntaxin 16 (S16528). For western blots, primary antibodies Anti-Syntaxin 16 monoclonal rabbit antibody EPR9156 (Abcam, ab134945), and Actin Loading Control mouse monoclonal Antibody BA3R (Invitrogen, MA5-15739) were used. Secondaries against mouse (Li-Cor IRDye 680RD Goat anti-Mouse IgG) and against rabbit (Li-Cor IRDye 800CW Goat anti-Rabbit IgG) were used.

### Enzyme-linked Immunosorbent spot (ELISpot) assays

ELISpot MSHA 96-well plates (Millipore) were coated with an anti-IFN-γ (Mabtech; #1-D1K) overnight at 4°C. After washing with PBS, plates were blocked with RPMI (Gibco) + 10% human serum + 2% L-Glutamine (Gibco) + 0.1% gentamycin (Gibco) for at least 1h at RT. BEAS-2B that were lipofected were seeded in a 6-well plate (Corning). After 24 hours, the cells were harvested, counted, re-plated, and allowed to adhere for at least 4h at 37°C. Cells were then infected with an MOI of 8 based on titer plates for the frozen stock. Infected antigen presenting cells were harvested after overnight incubation using RPMI + human serum and resuspended at 2e5 cells per mL; serial dilutions were then performed to generate 1:2, 1:4, and 1:8 dilutions. The cell dilutions were plated at 100μL/well in the MSHA plates in duplicate. PHA was used as a positive control for all ELISpot assays (1μg/well). After 1 hour, T cells were added at 1e4 cells/well (50μL/well) resulting in a final volume of 150μL/well. After overnight incubation at 37°C and 5% CO_2_, plates were washed in PBS + Tween, incubated with alkaline phosphatase-coupled secondary antibody (Mabtech; #7-B6-1-ALP) for 2 hours at RT, washed in PBS + Tween, washed in PBS, then incubated with BCIP/NBT substrate (Mabtech) for 10 minutes at RT. Substrate was washed off with de-ionized H_2_O and dried before measuring IFN-γ spot forming units (SFU) on an AID ELISpot reader. The mean of technical replicates was used to pool data from different experiments.

### Flow cytometry

BEAS-2B constitutively over-expressing MR1-GFP under a minimal CMV promoter were seeded in a 6-well plate (Corning) and lipofected. The next day, the cells were treated with 100 µM 6-FP (Schirks Laboratories) or the equivalent volume of 0.01M NaOH (Sigma). The next day, the cells were harvested, washed in flow buffer (PBS + 2% goat serum, 2% human serum, 0.5% FBS) and then stained with APC-conjugated mouse IgG2a, κ Isotype Control antibody (clone MOPC-173; BioLegend, #400220), APC-conjugated anti-MR1 antibody (clone 26.5; BioLegend, #361108), or APC-conjugated anti-human HLA-A,B,C antibody (clone W6/32; BioLegend, #311410) at 4°C for at least 30 min. Cells were then washed in PBS and fixed in 1% paraformaldehyde (Electron Microscopy Sciences) for at least 20 minutes at room temperature. Samples were acquired on a BD LSR II cytometer at the OHSU Flow Cytometry Core, and data were analyzed using FlowJo v10.8.1. Software (BD Life Sciences). Cells were gated based on forward scatter (FSC) area and side scatter (SSC) area, followed by two Single Cell gates based on 1) FSC width vs FSC height and 2) SSC width vs SSC height.

### Microscopy

Missense control and knockdown BEAS-2B:TET-MR1GFP LV cells were plated in 8-well Nunc #1.5 slides without doxycycline treatment. The cells were fixed in 4% PFA for 30 minutes. Next, the cells were permeabilized with flow buffer containing 0.2% saponin for 30 minutes. Staining was conducted with golgin-97 and Syntaxin 16 antibodies at 1:100 for 45 minutes. Goat-anti-mouse Alexa 568 (Invitrogen) and Goat-anti-rabbit Alexa 647 (Invitrogen) were used for secondary antibodies at 1:1000 for 45 minutes. Finally, the cells were stained with DAPI and Fluoromount G mounting media(SouthernBiotech) was added. The cells were imaged on a DeltaVision Wide field Deconvolution microscope with a 60x objective and a Nikon Coolsnap ES2 HQ. Each image was acquired as Z-stacks in a 1024×1024 format. A total of 11 images with 18 cells were imaged for the Missense control group, and 14 images with 30 cells were imaged for the Stx16 knockdown group across two independent experiments. Images were processed on Imaris (Bitplane) using “Surface” function to generate surfaces of Stx16 and golgin-97. The sum of the fluorescence intensity for the specific channel per cell was plotted.

For live-cell imaging, BEAS-2B:TET-MR1GFP LV cells were transiently transfected with 1.5 ug plasmid per 1.0e6 cells using an Amaxa Nucleofector (Kit T, program G-016). After transfection, the cells were plated into 4-well 1.5 mm glass bottom chamber slides (Nunc). Doxycycline was added at 1.0 µL/mL (2.0 mg/mL stock). The cells were imaged the next day on a DeltaVision Wide field Deconvolution microscope with a 60x objective and a Nikon Coolsnap ES2 HQ. Hoechst was added 30 minutes before imaging. Each image was acquired as Z-stacks in a 1024×1024 format. Images were processed on Imaris (Bitplane) and co-localization was analyzed using “Spots” function setting a distance threshold of 0.5 µm. A total of 36 images were acquired for Stx12-RFP across two independent experiments, and 26 images for Stx16-RFP were acquired across three independent experiments.

### Knockdown of Candidate Gene Expression

BEAS-2B plated in 6-well tissue culture plates (Corning) at 70% confluency were transfected with 50nM Silencer Select siRNA using HiPerFect (Qiagen) or Lipofectamine RNAimax Transfection Reagent (Invitrogen). Cells were incubated at 37°C for 48 hours. For functional assays, the knockdown cells were infected with Mtb and used the following day in an IFN-γ ELISpot assay, plated for microscopy, or treated with 6-FP and the following day processed and stained for Flow Cytometry.

### RT-qPCR analysis

Total RNA was extracted and isolated from BEAS-2Bs using the RNeasy Mini Kit (Qiagen). RNA was converted to cDNA using the High-Capacity cDNA Reverse Transcription Kit (Life Technologies). cDNA was purified using the PCR Purification Kit (Qiagen). RT-qPCR was performed using TaqMan Universal PCR Master Mix (Thermo Fisher Scientific) and Taqman FAM-MGB Gene expression Assay probes (Thermo Fisher Scientific) to detect transcripts of Stx6 (Hs01057343_m1), Stx12 (Hs00295291_m1), Stx16 (Hs01002372_m1), MR1 (Hs00155420_m1), and GAPDH (Hs02758991_g1) as an internal reference. Analysis was performed on a Step One Plus Real-Time PCR System (Applied Biosystems). Expression levels normalized to *Gapdh* (ΔC_T_) and relative expression levels normalized to *Gapdh* (2^−ΔΔCT^) were calculated.

### Western Blot

BEAS-2B:TET-MR1GFP LV cells were resuspended in 4x Laemmli Sample Buffer (Bio Rad) with 10% 2-Mercaptoethanol. Cells were incubated at 95°C for 5 minutes before loading onto a 4-20% polyacrylamide gel (Bio Rad). Gels were run at 95V for about 1.5 hours and transferred onto a polyvinylidene fluoride membrane (Millipore), which was then blocked in Odessey blocking buffer (Li-Cor) for one hour at room temperature. Membranes were then incubated with primary antibodies against Stx16 and loading control beta-actin on a shaker at 4°C overnight. Membranes were washed repeatedly in PBS + 0.1% Tween-20 and incubated with secondary antibodies against mouse and rabbit conjugated to different IRDyes. Blots were imaged on a Li-Cor Odyssey imaging system.

### Data analysis and Statistics

Data were analyzed with GraphPad Prism 10.2. Figures were assembled in Inkscape. Statistical significance for ELISpot assays was determined by nonlinear regression using agonist versus response (3 parameters) to compare differences in best-fit values of top and EC_50_ between the curves. Top and EC_50_ values were determined by least-squares regression, and the fit of one curve fit to all the data sets was compared with the fit of individual curves fit to each data set using the extra sum-of-squares F test. Statistical significance for microscopy was determined using Mann-Whitney test for nonparametric data. Statistical significance for flow cytometry was determined using two-tailed paired t-test.

## Acknowledgements

We thank Corinna Kulicke and Megan Huber, Oregon Health and Science University, for insightful discussions. We thank Wilmon Grant for construction of the Stx12-RFP plasmid. We acknowledge expert technical assistance by staff in the Advanced Light Microscopy Core in the Department of Neurology and Jungers Center at Oregon Health and Science University, and the assistance of the Oregon Clinical & Translational Research Institute, which is supported by the National Center for Advancing Translational Sciences, National Institutes of Health, through Grant Award Number UL1TR002369. This project used the OHSU Flow Cytometry and Monoclonal Antibody Shared Resource Core Facility (RRID:SCR_009974) for analytical flow cytometry. We would like to thank the participants who gave time and dedication to this health research. This work was supported by NIH grants AI134790 (DML), AI151079 (EK) and AI153359 (EK), AI186352-01A1 (AT), AI170496-02 (AT), and GM142619-01 (AT). This work was also supported in part by Merit Award #I01 BX000533 from the U.S. Department of Veterans Affairs Biomedical Laboratory (DML). The contents do not represent the views of the U.S. Department of Veterans Affairs or the United States Government.

## Author Contributions

The experiments presented were conceptualized by EK, AT, and DML. EK, AT, JCP, and AW performed experiments and analyzed data. EK and DML supervised the work and provided advice and technical expertise. All authors contributed to revising and reviewing the manuscript. All authors approved the final version of the manuscript.

## Data Availability Statement

All data generated and analyzed during this study are included in this published article.

