## Supplementary Material for "Sorting endosomes play key roles in presentation of *Mycobacterium tuberculosis*-derived ligands to MAIT cells"

**Title**

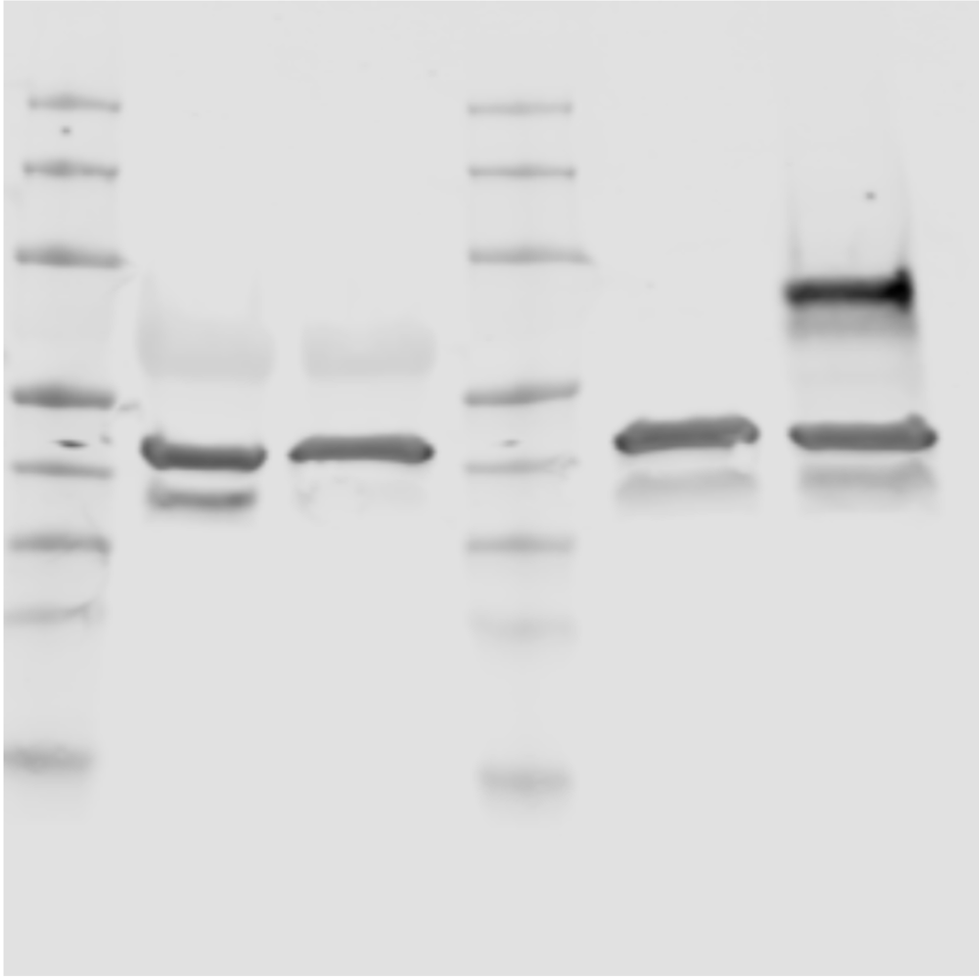

**Supplementary Figure S1: Full-length blot of Figure 3A.** From left to right, lanes 1-3 are from a separate experiment not included in Figure 3A. Lane 4 contains ladder, lane 5 contains non-target control transfected BEAS-2B:TET-MR1GFPs, and lane 6 contains Stx16-RFP transfected BEAS-2B:TET-MR1GFPs.

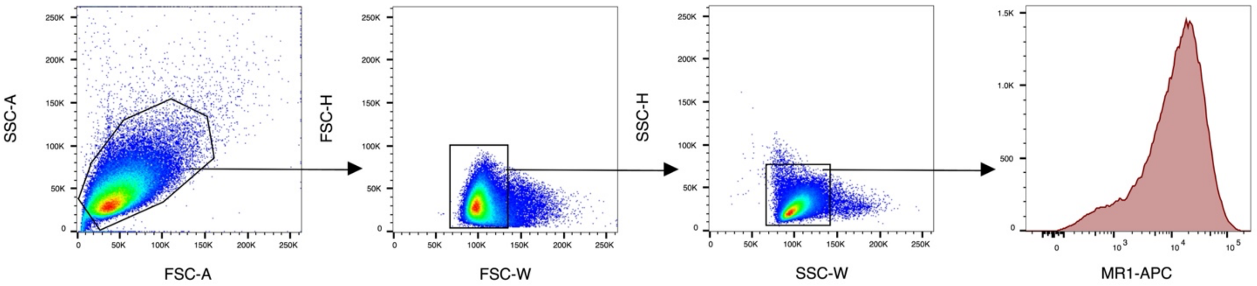


**Supplementary Figure S2: Gating strategy for MR1 surface staining.**
